# Voltage-Dependent Ca^2+^ Release Is Impaired in Hypokalemic Periodic Paralysis Caused by Ca_V_1.1-R528H but not by Na_V_1.4-R669H

**DOI:** 10.1101/2022.05.17.492380

**Authors:** Marino DiFranco, Stephen C. Cannon

## Abstract

Hypokalemic periodic paralysis (HypoPP) is a channelopathy of skeletal muscle caused by missense mutations in the voltage sensor domains (usually at an arginine of the S4 segment) of the Ca_V_1.1 calcium channel or of the Na_V_1.4 sodium channel. The primary clinical manifestation is recurrent attacks of weakness, resulting from impaired excitability of anomalously depolarized fibers containing leaky mutant channels. While the ictal loss of fiber excitability is sufficient to explain the acute episodes of weakness, a deleterious change in voltage sensor function for Ca_V_1.1 mutant channels may also compromise excitation-contraction coupling (EC-coupling).

We used the low-affinity Ca^2+^ indicator OGN-5 to assess voltage-dependent Ca^2+^-release as a measure of EC-coupling for our knock-in mutant mouse models of HypoPP. The peak Δ*F/F*_0_ in fibers isolated from Ca_V_1.1-R528H mice was about two-thirds of the amplitude observed in WT mice; whereas in HypoPP fibers from Na_V_1.4-R669H mice the Δ*F/F*_0_ was indistinguishable from WT. No difference in the voltage dependence of Δ*F/F*_0_ from WT was observed for fibers from either HypoPP mouse model. Because late-onset permanent muscle weakness is more severe for Ca_V_1.1-associated HypoPP than for Na_V_1.4, we propose the reduced Ca^2+^-release for Ca_V_1.1-R528H mutant channels may increase the susceptibility to fixed myopathic weakness. In contrast the episodes of transient weakness are similar for Ca_V_1.1- and Na_V_1.4-associated HypoPP, consistent with the notion that acute attacks of weakness are primarily caused by leaky channels and are not a consequence of reduced Ca^2+^-release.

## INTRODUCTION

Hypokalemic periodic paralysis (HypoPP) is a dominantly inherited disorder of skeletal muscle that presents with recurrent episodes of weakness lasting for hours to a day or even longer (1, 2). Recovery of strength is initially complete, with normal muscle function between attacks, but mild to moderate permanent weakness and vacuolar changes in muscle may occur with aging. HypoPP is an ion channelopathy caused by missense mutations of *CACNA1S* encoding the Ca_V_1.1 subunit of the L-type calcium channel (3), or less frequently by missense mutations of *SCN4A* encoding the Na_V_1.4 subunit of the sodium voltage-gated channel (4). Remarkably, 24 of the 25 reported HypoPP mutations are missense substitutions at arginine residues in S4 transmembrane segments of the voltage-sensor domains of these channels (5, 6).

The transient episodes of weakness are caused by sustained depolarization of the fiber resting potential (*V_rest_*) that inactivates sodium channels and thereby impairs fiber excitability (1). This depolarized shift of *V_rest_* paradoxically occurs in low extracellular [K^+^] (< 3.5 mM) because of the anomalous “gating pore” leakage current created by the missense mutations of S4 in either Ca_V_1.1 or Na_V_1.4 (7–9). Because the S4 segment is a critical component of the Ca_V_1.1 voltage sensor that couples transverse tubule depolarization to activation of the ryanodine receptor and Ca^2+^ release from the sarcoplasmic reticulum (SR), the possibility of a defect in excitation-contraction coupling (EC-coupling) has been proposed for the pathogenesis of HypoPP.

The integrity of EC-coupling in Ca_V_1.1-associated HypoPP has been assessed in only two prior studies (9, 10), which had conflicting results, and has never been examined for Na_V_1.4-associated HypoPP. Jurkat-Rott and colleagues (10) did not detect a difference for Ca^2+^-release between WT and Ca_V_1.1-R528H muscle, using fura-2 microfluorimetry in voltage-clamped human HypoPP fibers (WT/R528H heterozygous) or in a mouse muscle cell line that is null for Ca_V_1.1 (GLT cells) with transiently expressed R528H mutant channels. We created a knock-in mutant mouse model of Ca_V_1.1-R528H with a robust HypoPP phenotype and measured voltage-dependent Ca^2+^ transients with fluo-4 in dissociated fibers (9). Ca^2+^-release was comparable for WT and heterozygous Ca_V_1.1-R528H fibers, but the average response from a pool of 14 homozygous Ca_V_1.1-R528H fibers was markedly reduced to 30% of WT levels.

In the present study, we use a low-affinity fast Ca^2+^ dye, OGN-5, to address the prior conflicting results for Ca_V_1.1-R528H (9, 10), and also assess Ca^2+^-release for Na_V_1.4-R669H fibers. We confirm a reduced peak Δ*F/F*_0_ for Ca_V_1.1-R528H fibers, but at a more modest degree of about two-thirds of WT levels. We propose this milder impairment of Ca^2+^-release compared to our prior report is because more stringent criteria were used to select healthy fibers that presumably do not have the triad disruption that occurs with vacuolar myopathy in HypoPP. No defect of Ca^2+^-release was detected for Na_V_1.4-R669H fibers.

## MATERIALS AND METHODS

### Mouse Models of HypoPP

The generation of knock-in mutant mouse models for hypokalemic periodic paralysis (HypoPP), screening progeny for mutant alleles, confirmation of mutant allele expression, and characterization of the periodic paralysis phenotype with paradoxical depolarization of *V_rest_* and loss of force in a low-K^+^ challenge has been previously described (9, 11). Two murine models of HypoPP were studied here: a missense mutation of *Cacna1s* coding for Ca_V_1.1 (mR528H) and a missense mutation of *Scn4a* coding for Na_V_1.4 (mR663H). These homologous mutant alleles in mice correspond to the human disease-causing mutations Ca_V_1.1-R528H and Na_V_1.4-R669H, and this nomenclature is used herein to facilitate the comparison to the clinical literature. Mouse lines have been propagated in the 129/sv strain for more than 20 generations. All procedures on mice were in accordance with animal protocols approved by the Institutional Animal Care and Use Committee at the David Geffen School of Medicine, University of California, Los Angeles, CA.

### Muscle Fiber Preparation

All recordings were performed on dissociated single fibers from the flexor digitorum brevis muscle (FDB) as previously described (12). Animals aged 2 – 4 months were euthanized by isoflurane inhalation followed by cervical dislocation. Both male and female mice were used, and the data were pooled. The FDB was rapidly dissected free and pinned to the bottom of a sylgard-coated dish containing collagenase type II (6 mg/ml, Gibco) in standard Tyrode’s solution. The dish was gently agitated for 45 min in an incubator at 35 ^o^C. The enzymatically digested muscle was then triturated in a series of fire-polished glass pipettes, with progressively smaller tip diameters. Dissociated single fibers were rinsed 4 times with collagenase-free Tyrode’s solution and then transferred to a glass-bottomed recording chamber constructed from a 35 mm plastic dish fitted with a glass coverslip (FisherFinest #12-548-A).

### Electrophysiology

Single FDB fibers of length 400 to 600 μm (mean 537 μm) were impaled with two sharp microelectrodes (10-12 MΩ), near the midpoint and longitudinally separated by 10-15 μm. Recordings were initiated in two-electrode current-clamp (TEV-200, Dagan Corporation), and a holding current was applied to polarize the fiber to −80 mV. We intentionally selected “healthy” fibers, as defined: (i) optically by sharp edges, clear sharply contrasted striations, and no intracellular inclusions (ii) electrophysiologically by a spontaneous membrane potential of −40 mV to −50 mV upon initial impalement, and full polarization to −80 mV with a holding current in the range of −5 to −25 nA in Tyrode’s solution.

After the fiber equilibrated for 10 minutes at −80 mV, action potentials were elicited by application of a brief depolarizing current of 0.2 to 0.3 μA for 0.5 msec. Fiber contraction was prevented by using an internal solution for the microelectrodes that contained a high concentration of EGTA (see Solutions below).

To improve the voltage control of the clamped fiber (speed, spatial uniformity, steady-state error), we exchanged the Tyrode bath with a TEA-Cl solution that also contained blockers for sodium, calcium, and chloride channels (see Solutions below). The holding current was manually adjusted to keep *V_m_* near −80 mV, and then the TEV-200 amplifier was switched to voltage-clamp mode at a clamp potential of −80 mV. The voltage protocol was a sequence of 10 msec step depolarizations, over a range from −80 mV to +80 mV, from a holding potential of −80 mV. The sequence progressed from −80 mV to more positive values in 10 mV increments, and the fiber was returned to −80 mV for 20 sec between pulses. Current (low-pass filtered at 5 KHz) and voltage signals were sampled at 25 KHz using a 16-bit A/D and D/A converter, controlled with Labview software (National Instruments).

### Detection of Ca^2+^ transients

A fluorescent, low affinity, impermeant Ca^2+^-sensitive dye, Oregon Green 488 BAPTA-5N (OGN-5), was used to follow fast changes of myoplasmic [Ca], in response to action potentials or voltage pulses. The dye was excited at 480 +/-20nm, and its emission was restricted to 535 +/-22nm. Excitation and emission bandwidths were separated by a 510 dichroic mirror. All optical components were from Semrock. The light source was an LED (SP-05-B6, Luxeon StarLeds) driven by a custom-made power source under computer control. The experimental chamber was placed on the stage of an inverted microscope (Olympus XI-171) equipped with a homemade epifluorescence attachment. Fibers were imaged with a 100x, 1.4NA oil immersion objective. The illumination spot diameter was adjusted to about 90% of the fiber diameter. The light detector consisted of a low capacitance PIN photodiode (UV-001, OSI Optoelectronics) connected to the head stage of a patch-clamp amplifier (Axopatch 2B, Molecular Devices Inc.) with no bias current applied. The dark current was set to zero. The photocurrent was filtered at 2 kHz. Changes in myoplasmic [Ca] are presented as Ca^2+^-dependent fluorescence transients, *F*, expressed as fractional changes above the base line, Δ*F/F*_0_ = (*F* − *F*_0_)/*F*_0_, where *F*_0_ is the pre-stimulus photocurrent.

### Solutions

The standard Tyrode’s solution contained (mM): 150 NaCl, 4 KCl, 2 CaCl_2_, 1 MgCl_2_, 10 glucose, and 10 MOPS, pH 7.4 with NaOH. The extracellular solution for voltage-clamp studies contained (mM): 140 TEA-Cl, 10 CsOH, 2 CaCl_2_, 1 MgCl_2_, 10 glucose, 10 MOPS, pH 7.4 with HCl. Channel blockers were added from stock solutions for a working concentration of 200 nM TTX, 20 μM nifedipine, 200 μM 9-anthracene carboxylic acid. The internal solution contained (mM): 140 K-aspartate, 20 K-MOPS, 5 MgCl_2_, 5 Na_2_ creatine phosphate, 5 ATP-K_2_, 5 glucose, 5 reduced glutathione, 30 EGTA and 15 mM Ca(OH)_2_ for a 30:15 EGTA:Ca^2+^ ratio equivalent to 80 nM free Ca^2+^. The membrane impermeant Ca^2+^ indicator OGB-5N (ThermoFisher) was added to the internal solution for a final concentration of 125 μM. All experiments were performed at room temperature (22 °C).

### Statistical Analysis

Data are presented as mean ± SEM in Tables or error bars in Figures. ANOVA was used to test for statistically different mean values (*p* < 0.05) amongst the 4 genotypes (WT, heterozygous Ca_V_1.1-R528H, homozygous Ca_V_1.1-R528H, homozygous Na_V_1.4-R669H).

## RESULTS

### Ca^2+^ Release by an Action Potential is Reduced for Ca_V_1.1-R528H but Not for Na_V_1.4-R669H HypoPP Fibers

The resting potential of acutely dissociated FDB fibers is typically −40 mV to −50 mV (12), a range that is substantially depolarized from the value of −80 mV to −90 mV measured from intact whole muscle using sharp microelectrodes (13). Therefore, we applied a holding current to set *V_m_* = −80 mV after fiber impalement with two microelectrodes. This paradigm also compensates for the mild depolarization of *V_rest_* in HypoPP fibers (9, 11), so that action potentials were elicited from the same initial conditions for fibers from WT and mutant mice. Representative action potentials triggered by a 0.5 msec current pulse of 0.2 to 0.3 μA, and the corresponding ΔF/F for the Ca^2+^transient, are shown for WT and homozygous Ca_V_1.1-R528H fibers in Figure 1. The ΔF/F waveforms were similar, but the peak amplitude was consistently smaller for homozygous Ca_V_1.1 - R528H fibers compared to WT. The distribution of peak Δ*F/F*_0_ values recorded in FDB fibers from each of the mouse lines is shown in the box plot of Figure 2. On average, the peak Δ*F/F*_0_ for homozygous Ca_V_1.1-R528H was 66% of WT (0.48 ± 0.038, n = 12 and 0.73 ± 0.028, n = 18; respectively, *p* < 0.0001 ANOVA). Heterozygous WT / Ca_V_1.1-R528H fibers had an intermediate peak Δ*F/F*_0_ = 0.62 ± 0.047, n = 10 that was not statistically different from WT. The peak Δ*F/F*_0_ for homozygous Na_V_1.4-R669H fibers (0.72 ± 0.074, n = 8) was identical to that of WT.

**Figure 1.**
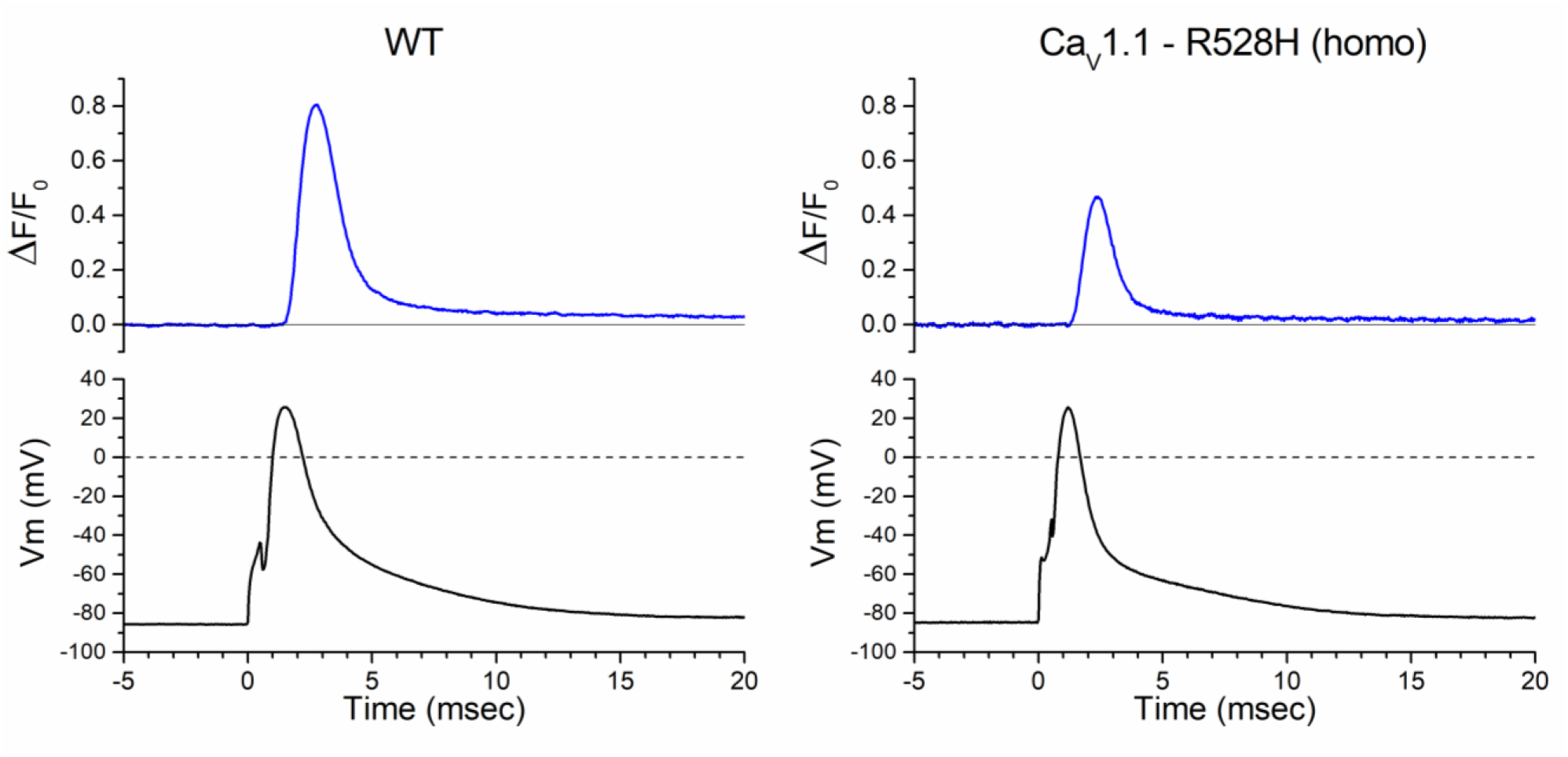
Ca^2+^ release elicited by an action potential. Each record shows the Δ*F/F*_0_ response for a single trial, in which an action potential was triggered in an isolated FDB fiber by a 0.5 msec current pulse. These representative traces show a larger Δ*F/F*_0_ for WT (*left*) compared to a fiber from a homozygous Ca_V_1.1-R528H mouse (*right*).

**Figure 2.**
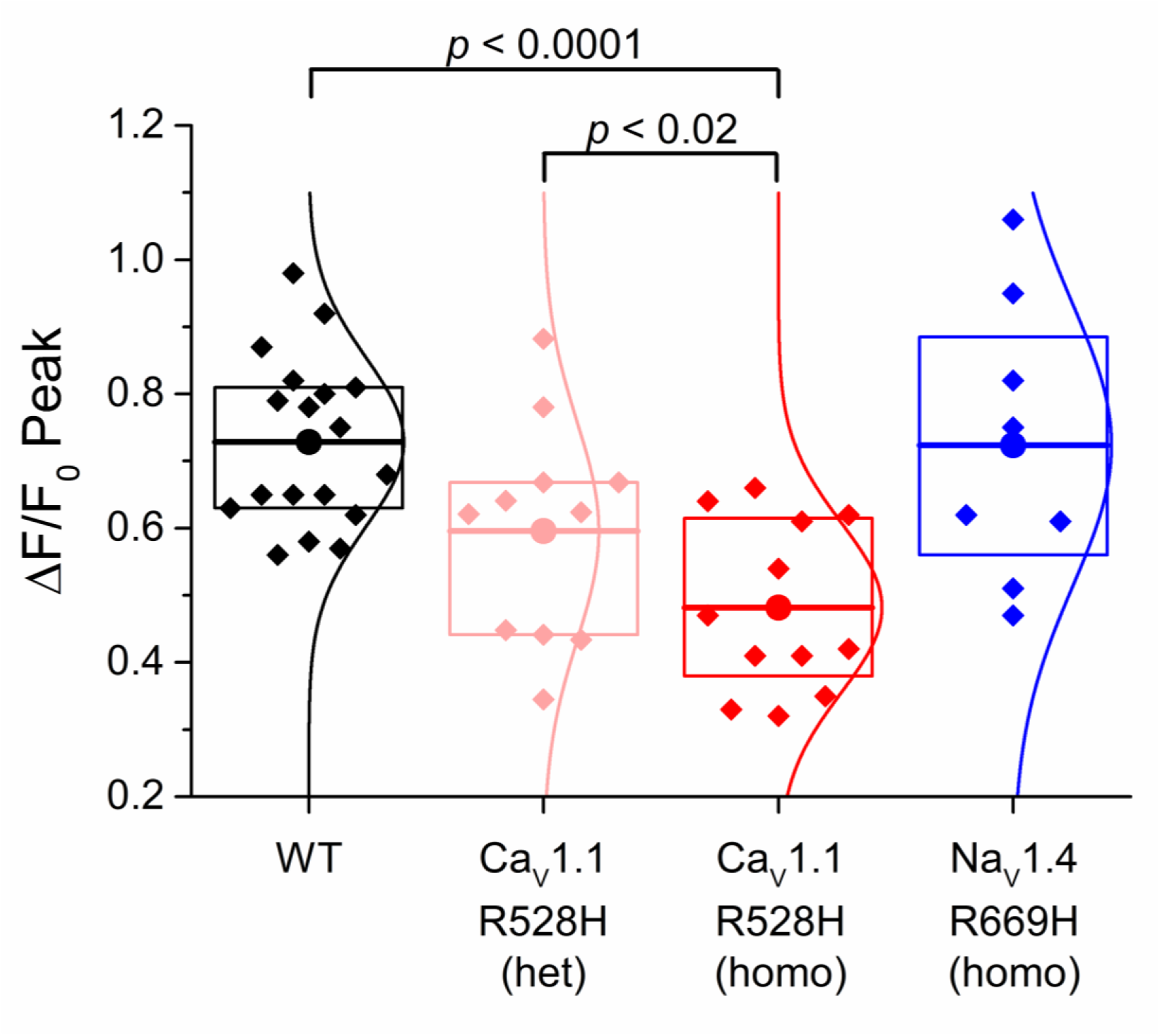
Ca^2+^ release by an action potential is smaller for Ca_V_1.1-R528H fibers than for WT. Data points in the box plots show the peak Δ*F/F*_0_ recorded for individual FDB fibers. The sample mean (large circle and horizontal line) was lower for homozygous Ca_V_1.1-R528H than WT (*p* < 0.001) or for heterozygous Ca_V_1.1-R528H (*p* < 0.02); whereas the peak Δ*F/F*_0_ for homozygous Na_V_1.4-R669H fibers was not different from WT. Box dimensions show 25^th^ and 75 percentiles, and the smooth curve is a Gaussian best fit.

To determine whether the smaller peak Δ*F/F*_0_ for homozygous Ca_V_1.1-R528H fibers was possibly a consequence of a smaller amplitude action potential in HypoPP fibers, we generated a scatter plot of Δ*F/F*_0_ as a function of the ΔVm for the action potential for each fiber (Figure 3). In the overlapping range of action potential amplitudes (100 mV to 115 mV), the peak Δ*F/F*_0_ was consistently smaller for homozygous Ca_V_1.1-R528H fibers than for WT ones. A linear fit to those data show that Ca^2+^ release has a comparable voltage sensitivity for WT and homozygous Ca_V_1.1-R528H (despite a partial loss of charge in the DII voltage sensor), smaller Δ*F/F*_0_ with smaller ΔV, but the response for the latter is shifted downward to lower values.

**Figure 3.**
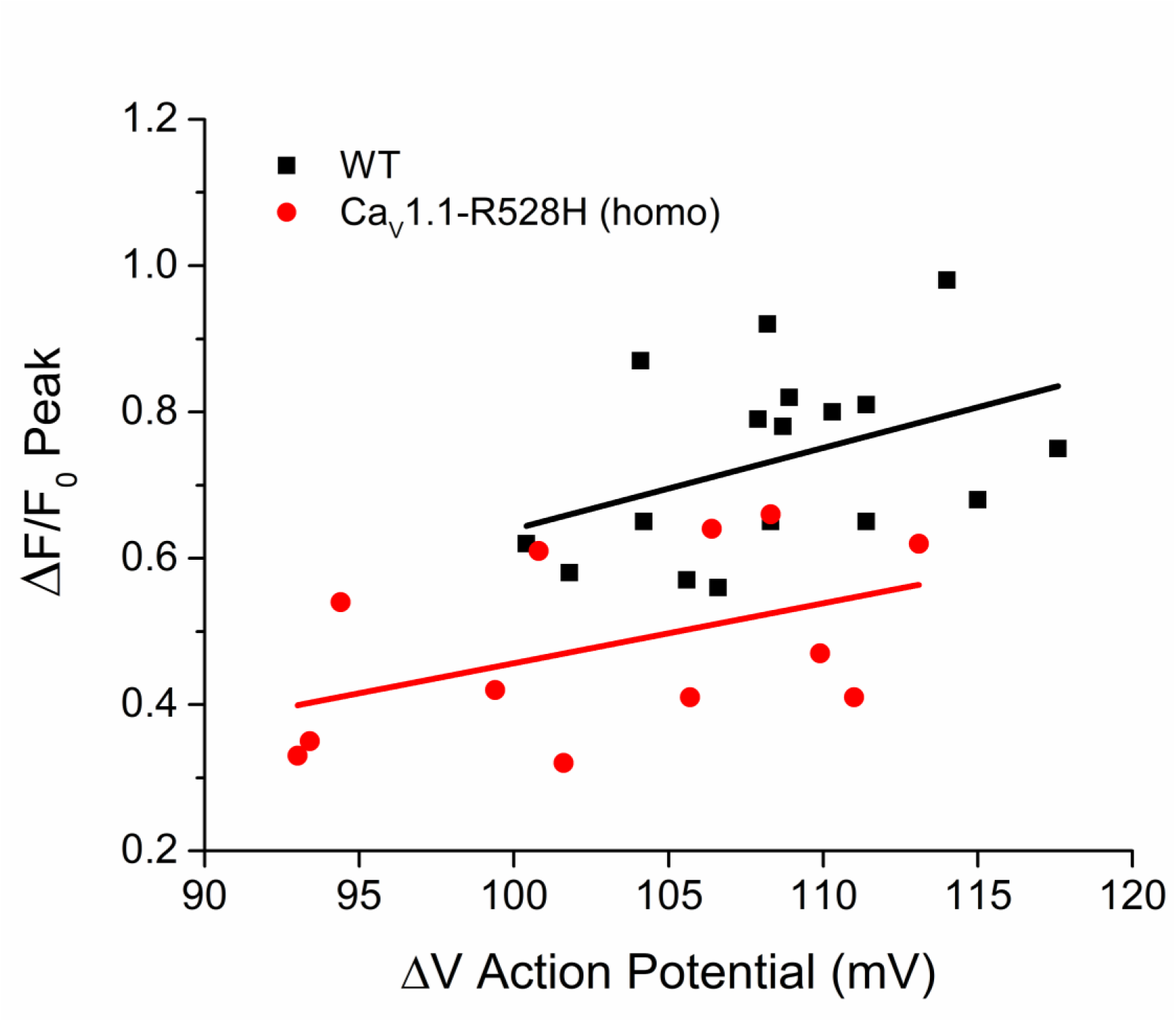
Reduced Ca^2+^ release for Ca_V_1.1-R528H fibers is not because the action potential has a smaller amplitude. The Δ*F/F*_0_ is plotted as a function of ΔV for the corresponding action potential. Each point represents the response of a single trial from a different fiber. Lines show the least-squares fit for all the data from WT (black squared) or homozygous Ca_V_1.1-R528H (red circles).

We previously reported a modest disruption of the coupling between voltage-sensor movement and channel opening for Na_V_1.4-R669H (14). This gating defect may result in a reduction of the peak Na^+^ current, and consequently smaller amplitude action potentials. Consistent with this prediction, Δ*V_m_* for action potentials elicited in Na_V_1.4-R669H fibers were smaller than for WT fiber (102 ± 2.3 mV n = 6; 108 ± 1.2 mV n = 13; respectively *p* < 0.05). However, this small reduction in Δ*V_m_* for the action potential was not associated with a reduction of the peak ΔF/F for Na_V_1.4-R669H (Figure 2). Amplitudes for action potentials in Ca_V_1.1-R528H fibers (homozygous and heterozygous) were not different from those in WT.

### A Larger Holding Current is Required for HypoPP Fibers, Consistent with a Gating Pore Leakage Current

The primary functional defect in HypoPP is the anomalous gating pore leak current, resulting from missense mutations for the S4 segment in voltage sensor domains of Ca_V_1.1 (9, 15) or Na_V_ 1.4 (7, 8). The gating pore leak, although pathogenic, is only about 2% of the resting fiber conductance (9, 11, 15, 16), and so the detection of this anomalous current by voltage-clamp is technically challenging. Consequently, the voltage-clamp strategy requires either blocking all of the ion channels (conventional pores) in a HypoPP muscle fiber or using the oocyte system to achieve high expression of the mutant channel. The current-clamp studies herein, provide a new opportunity to detect the presence and magnitude of an anomalous inward current in HypoPP fibers, consistent with the gating pore current. Importantly, this measurement was performed in a standard mammalian bath solution without ion channel blockers. The box plot in Figure 4 shows the distribution of holding currents that were required to set *Vm*= −80 mV. On average, larger negative holding currents were required to hyperpolarize HypoPP fibers, homozygous for either Ca_V_1.1-R528H or Na_V_1.4-R669H. As expected for heterozygous Ca_V_1.1-R528H with half the gating pore current density, an intermediate holding current was required, between WT and homozygous HypoPP. Moreover, the magnitude of the additional negative holding current (about −10 μA/cm^2^) is consistent with the reported conductance (10 to 20 μS/cm^2^) and reversal potential (−20 mV) for the anomalous gating pore current in murine HypoPP fibers (9, 11).

**Figure 4.**
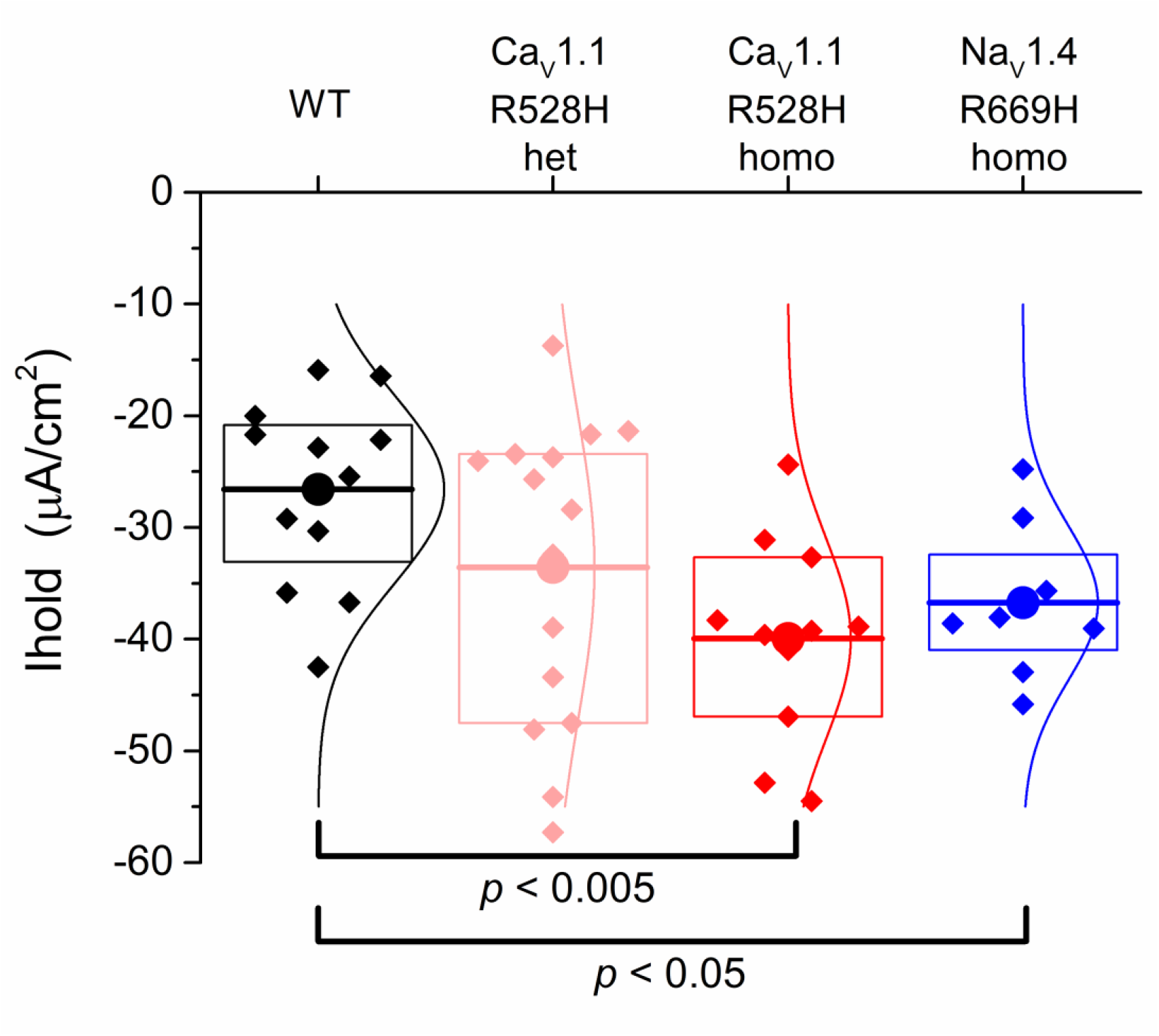
The holding current required to maintain the fiber *V_m_* = −80 mV is larger (more negative) for HypoPP fibers than for WT. Each point in the box plot represents the original holding current required to set *V_m_* to −80 mV for an individual fiber. The response from a fiber was included in the dataset if the measured voltage several minutes later was in the range −88 to −78 mV. Large circles and horizontal lines show the mean, the box indicates the 25% and 75% range, and the smoother curves are the Gaussian fit.

### Voltage Dependence of Ca^2+^ Release is not Altered by the HypoPP Mutations

The voltage dependence of Ca^2+^ release was assessed by measuring Δ*F/F*_0_ in voltage-clamped FDB fibers. Figure 5 shows set of responses observed for series of voltage steps from −40 mV to +80 mV, from a holding potential of −80 mV. The first detectable change in Δ*F/F*_0_ was at −30 mV (blue traces) for both WT and homozygous R528H fibers. The Δ*F/F*_0_ saturated for voltage steps > +30 mV, and this level was larger for WT than for homozygous Ca_V_1.1-R529H as shown by the representative traces in Figure 5. The peak Δ*F/F*_0_ is shown as a function of voltage in Figure 6A. The maximum Δ*F/F_0_,* determined by fitting a Boltzmann function to the responses for each fiber over the voltage range from −40 mV to +40 mV, was 76% of WT for homozygous Ca_V_1.1-R528H fibers (0.71 ± 0.026, n = 16 and 0.53 ± 0.028, n = 7 respectively; *p* = 0.01 ANOVA). The maximum Δ*F/F*_0_ for heterozygous Ca_V_1.1-R528H was 0.60 ± 0.062, n = 9, a value that lies between WT and homozygous Ca_V_1.1-R528H but is not statistically different from either (*p* > 0.05 ANOVA).

**Figure 5.**
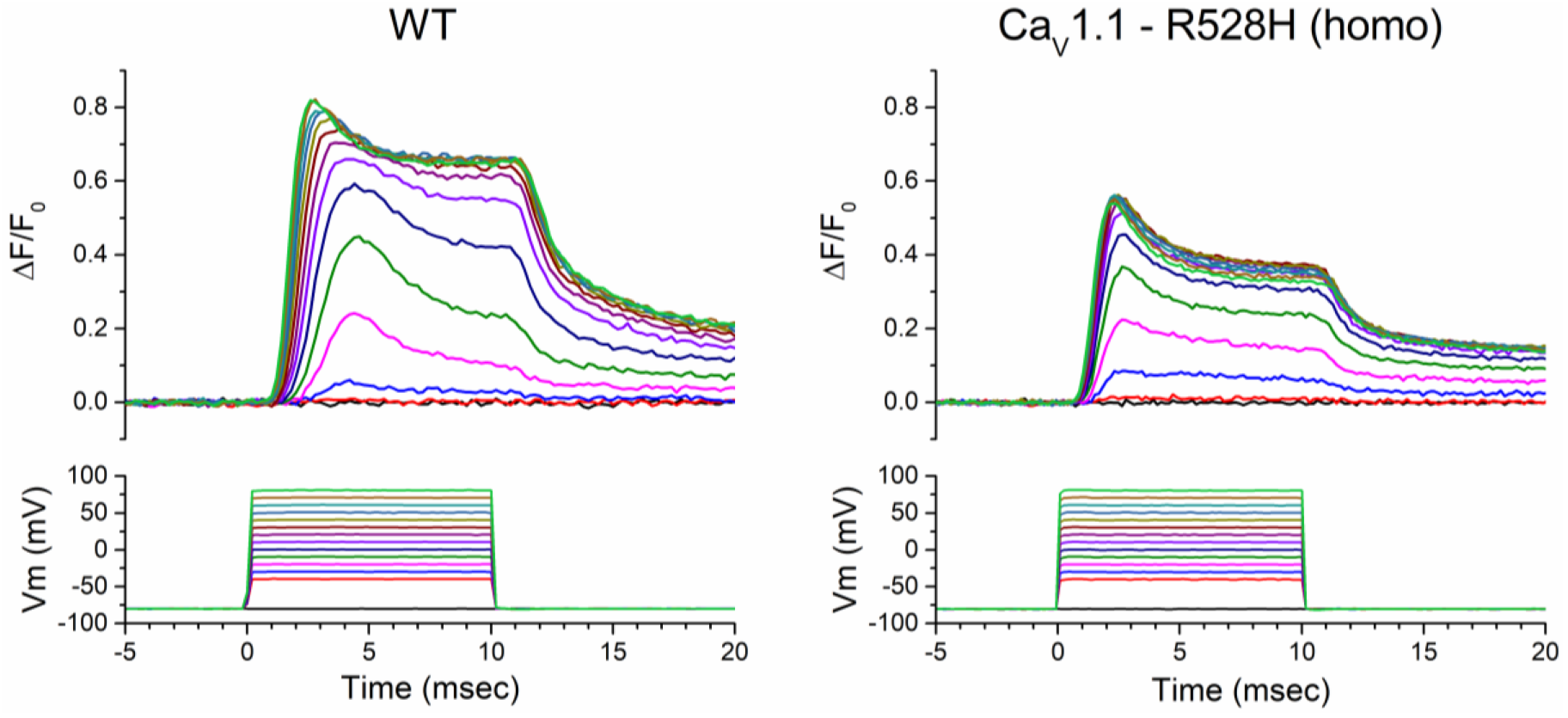
Ca^2+^ release transients in response to a series of step depolarizations for voltage-clamped FDB fibers. Each sweep is for a single trial (no averaging), and the responses are superimposed for voltage steps over the range −40 to +80 mV. Representative responses show the peak Δ*F/F*_0_ was larger for WT (*left*) than for homozygous Ca_V_1.1-R528H (*right).* The traces in the lower panels are the measured *V_m_*, not the digital command pulses.

**Figure 6.**
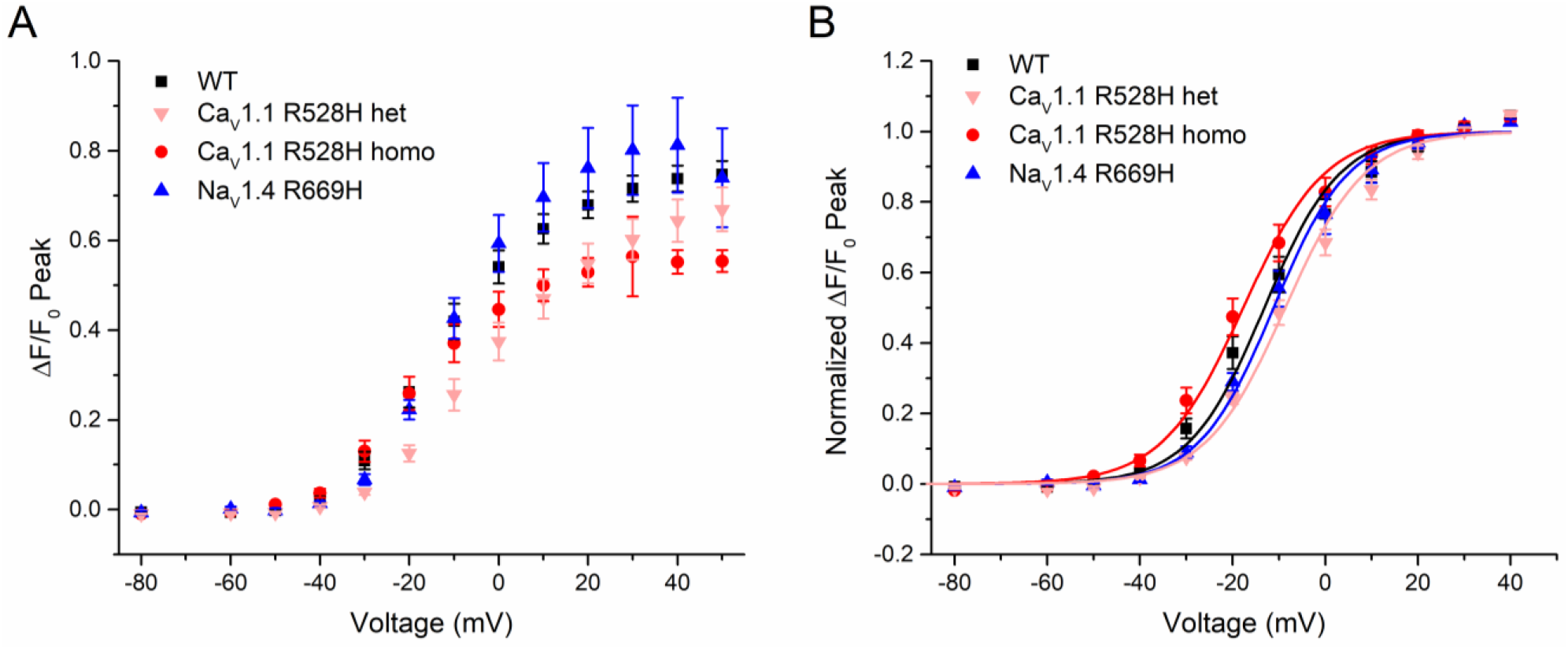
Voltage-dependent Ca^2+^ release is reduced for homozygous Ca_V_1.1-R528H fibers compared to WT fibers. (A) The peak Δ*F/F*_0_, averaged from measurements in several fibers, is shown as a function of step potential. (B) Amplitude-normalized Δ*F/F*_0_ shows a comparable voltage-dependence of Ca^2+^ release in fibers from WT and either HypoPP mutant line. Symbols show mean ± SEM.

A comparison for the voltage dependence of Δ*F/F*_0_ is illustrated with the amplitude-normalized data in Figure 6B. While some variability was apparent for the voltage of the midpoint, none of the differences in *V*_1/2_ amongst the four genotypes were statistically distinguishable. The steepness of the voltage dependence of Δ*F/F*_0_ was also identical, despite the partial loss of gating charge for Ca_V_1.1-R528H mutant channels. Parameter values for the Boltzmann fits (smooth lines, Figure 6B) are shown in Table 1.

**Table 1.** Parameters for Boltzmann fit of Δ*F*(*V*)/*F*_0_

| Genotype | $(\Delta F/F_0)_{max}$ | $V_{1/2}$ (mV) | $k$ (mV) | N |
| --- | --- | --- | --- | --- |
| WT | $0.71 \pm 0.026$ | $-13 \pm 2.7$ | $8.1 \pm 0.43$ | 16 |
| het Cav1.1-R528H | $0.60 \pm 0.061$ | $-8.2 \pm 1.4$ | $8.7 \pm 0.49$ | 9 |
| homo Cav1.1-R528H | $0.53 \pm 0.028^\dagger$ | $-18 \pm 2.3$ | $8.8 \pm 0.56$ | 7 |
| homo Nav1.4-R669H | $0.79 \pm 0.098$ | $-11 \pm 1.7$ | $8.1 \pm 0.92$ | 5 |

$^\dagger p = 0.01$

## DISCUSSION

The prevailing view for the pathogenesis of recurrent episodes of weakness in HypoPP is that an aberrant sustained depolarization of *V_rest_* inactivates Na^+^ channels, resulting in severe attenuation or even failure of action potential propagation (1). While this reduction of excitability that drives EC-coupling is sufficient to explain the impaired generation of contractile force, a possible contribution from altered voltage sensing or disrupted coupling to activation of RyR1 and Ca^2+^ release remains an open question (10, 17). The S4 segment in the domain II voltage sensor, containing R528H, was recently shown to translocate rapidly in response to membrane depolarization (*τ* = 3 msec) consistent with a role in EC-coupling, as opposed to the slow translocation of the domain I S4 segment (*τ* = 100 msec) that occurs with the same time course as channel opening (18). Indeed, the HypoPP mutations of *CACNA1S* are all located in voltage sensor domains (II-IV), and 9 out of 10 are missense substitutions at positively-charged arginine residues in S4 segments (19). This canonical pattern of HypoPP mutation sites has a common functional correlate. An anomalous “gating pore” leakage current, that can account for the paradoxical depolarization of *V_rest_* in low K^+^, has been reported for 7 HypoPP-Ca_V_1.1 mutations (16, 20–22). On the other hand, no gating pore current was detected for two mutations (R897K and R900S in IIIS4) which raises the possibility of other mechanisms for the intermittent weakness (21, 23).

The assessment of Ca^2+^-release integrity in HypoPP has previously been inadequate because of the challenges in obtaining muscle biopsies for this very rare disorder or for limited availability of suitable model systems. Only a single study of Ca^2+^-release in human HypoPP muscle has been reported (10), and these fura-2 measurements in voltage-clamped Ca_V_1.1-R528H fibers showed no difference in the midpoint or the steepness for the voltage-dependence of Ca^2+^-release. Amplitude-normalized responses were shown and no comment was made about the relative amplitude for HypoPP versus WT. In this same study, expression of Ca_V_1.1-R528H in Ca_V_1.1-null myotubes (GLT cells immortalized from the *mdg* mouse) also failed to reveal any differences in fura-2 transients compared to WT.

### The Reduced Ca^2+^-Release in Ca_V_1.1-R528H Fibers is Likely Because of Lower Functional Channel Density

Our measurements of Ca^2+^ transients using the low-affinity fast dye OGN-5 show a 24% to 34% reduction in the peak Δ*F/F*_0_ elicited by a voltage pulse or by an action potential, respectively, for homozygous Ca_V_1.1-R528H fibers. This value is comparable to the 36% reduction in maximal gating charge displacement, *Q_max_,* that we previously reported (9). Recordings from heterozygous Ca_V_1.1-R528H fibers show a milder reduction of peak Δ*F/F*_0_ (Figures 2 and 6) and of *Q_max_*(reference (9)), consistent with a dosage effect of the mutant allele. Taken together, these data suggest a reduced density of functional channels at the plasma membrane produces the reduction of peak Ca^2+^-release in Ca_V_1.1-R528H fibers. This interpretation also supports the notion that Ca_V_1.1-R528H channels are functionally equivalent to WT channels with regard to effectiveness for EC-coupling.

We previously reported a more pronounced reduction by 67% for the peak Δ*F/F*_0_ measured with Fluo-4 in homozygous Ca_V_1.1-R528H fibers in whole-cell recording mode (9). That study did not use the stringent criteria for fiber quality that were applied herein (see Materials and Methods). It is very likely that the population of fibers in our prior study included ones with structurally disrupted triads, as shown histologically by transverse tubular aggregates and vacuoles with dilated SR demonstrated on ultrastructural studies with transmission EM (9). We applied more stringent optical and electrical criteria for fiber integrity herein, in an effort to specifically assess the intrinsic capability of Ca_V_1.1-R528H channels for EC-coupling; rather than detecting the apparent effectiveness of EC-coupling in HypoPP muscle that may be compromised by a combination of altered Ca_V_1.1 function and structurally disrupted triads. The absence of a detectable impairment of Ca^2+^-release for Na_V_1.4-R669H also supports the notion that a primary disruption of EC-coupling is not a mechanism shared in common for HypoPP mutations in the pathogenesis of episodic weakness.

In contrast to our findings above, Jurkat-Rott and colleagues (10) concluded Ca^2+^-release “was not grossly altered” as reported by fura-2 measurements in human fibers heterozygous for Ca_V_1.1-R528H or for GLT cells expressing a uniform population of R528H HypoPP mutant channels. There are several plausible explanations for this discrepancy: (i) the reduction detected with the fast dyes (OGN-5, Fluo-4) may not be apparent with the slower high-affinity fura-2; (ii) the signal-to-noise was much less favorable in the fura-2 study, which was further exacerbated by the heterozygous state for the human fibers; (iii) the relative amplitude of Δ*F/F* for R528H mutant compared to WT was not reported in the fura-2 study, whereas both studies are concordant for no differences in *V*_1/2_ or *k* for the voltage dependence; (iv) the lower membrane expression of Ca_V_1.1-R528H that we detected in our HypoPP mouse model may not be a feature of human HypoPP muscle or of heterologous expression in GLT cells. Nevertheless, both studies agree that Ca_V_1.1-R528H does not disrupt the ability of the channel to participate in EC-coupling.

### Clinical Implications of Reduced Ca^2+^-Release in Ca_V_1.1-R528H HypoPP Muscle Fibers

The confluence of HypoPP mutations, almost all of which are in S4 segments of voltage sensors, raised questions about whether disrupted EC-coupling contributes to the acute attacks of weakness or to the late-onset permanent weakness. While a reduction of Ca^2+^-release is clearly present in our Ca_V_1.1-R528H mouse model of HypoPP, our overall impression is that a primary defect of EC-coupling is not a substantive contributing factor to the transient attacks of weakness. First, a “static” impairment of EC-coupling is more likely to act as a modifying effect for the severity of an episode of weakness rather than contributing to the occurrence of an ictal event. Second, the decrease in peak Ca^2+^-release after an action potential is only 24% in heterozygous Ca_V_1.1-R528H fibers, which is modest in comparison to the safety-factor of Ca^2+^ release for muscle contraction (24, 25). This interpretation is consistent with the clinical observation that muscle strength is normal between episodic attacks of HypoPP (1, 2). Third, the clinical severity of disease is equivalent for HypoPP from Na_V_1.4 mutations that are not expected to affect Ca^2+^-release, as now confirmed herein experimentally. Fourth, the electromyogram consistently shows a severe impairment of muscle excitability during an acute attack in HypoPP (26, 27), which is sufficient to produce the severe loss of force. The available data therefore support the view that the transient episodes of weakness in HypoPP are not a consequence of an intrinsic defect of EC-coupling.

The reduced Ca^2+^-release for Ca_V_1.1-associated HypoPP may contribute to the susceptibility to permanent muscle weakness with aging. In a cohort of 36 patients with HypoPP, permanent weakness was more pronounced for patients with Ca_V_1.1 mutations than with Na_V_1.4 mutations (15), consistent with the notion that impaired Ca^2+^-release contributes to permanent weakness. Moreover, studies in WT mice show an age-dependent decline in voltage-dependent Ca^2+^-release (50%) with an associated 35% decrease in maximal specific force of a tetanic contraction (28). These changes in WT muscle are caused by an age-dependent decrease of functional Ca_V_1.1 channels in the plasma membrane (29). We propose the reduced Ca^2+^-release for Ca_V_1.1-associated HypoPP and the decrease of functional Ca_V_1.1 channels with normal aging will synergistically increase the risk of permanent muscle weakness in HypoPP. This hypothesis also explains why patients with Na_V_1.4 mutations are less susceptible to permanent muscle weakness, because the channel defect does not reduce Ca^2+^-release. Thus far, studies of Ca^2+^-release in Ca_V_1.1-associated HypoPP are limited to R528H. Permanent muscle weakness is more severe and has an earlier age of onset for patients with the Ca_V_1.1-R1239H mutation (15). The prediction is Ca^2+^-release may be more severely impaired for R1239H than for R528H, which warrants additional studies of this and other Ca_V_1.1-associated HypoPP mutations.

## ACKNOWLEDGEMENTS

This work was supported by grants AR063182 and AR078198 from the National Institute of Arthritis, Musculoskeletal, and Skin Diseases of the NIH. The authors thank Marbella Quiñonez for maintaining the mouse colonies.

